# The role of asymmetric migration on local adaptation: an analysis of the classic two-habitat migration model

**DOI:** 10.1101/2025.01.30.635735

**Authors:** Tadeas Priklopil

## Abstract

We explore how migration and selection interact to shape local adaptation, revisiting the notion that increased gene flow typically suppresses genetic differentiation and local adaptation. Instead, we demonstrate that migration can both suppress and facilitate local adaptation, depending on the symmetry or asymmetry of migration rates between populations. This duality is examined in two scenarios within a single-locus theory: one in which local adaptation of a pair of alleles is maintained under arbitrary patterns of antagonistic selection, and another in which local adaptation develops through the recurrent invasion and fixation of alleles with weak selective effects on a continuous phenotype. For both scenarios, we analyze the classic two-habitat migration model, providing a novel examination of its dynamics. When migration is symmetric, an equal increase in migration rates impedes local adaptation by forcing both variant alleles to spend more time in non-local habitats where they are disfavored. In contrast, when migration rates increase asymmetrically, leading to skewed gene flow, both alleles spend more time in the same habitat, disadvantaging the non-local allele but favoring the locally adapted one. Thus, both increasing and decreasing migration rates can favor or hinder local adaptation, depending on the interplay between migration patterns and selection, underscoring the importance of migration heterogeneity in shaping local adaptation dynamics.

---

Local adaptation occurs when genetic polymorphisms are maintained, with each allele performing better in its native habitat than any immigrant allele from another habitat (Lenormand, 2002; Kawecki and Ebert, 2004; Blanquart et al., 2013). Gene flow, which mixes alleles between habitats, poses a key challenge to local adaptation by increasing the time alleles spend in habitats where they are at a selective disadvantage (Haldane, 1948; Slatkin, 1987; Lenormand, 2002; Yeaman, 2015; Hendry, 2017). In this note, we show that while this reasoning holds under symmetric migration, it breaks down under asymmetric migration. When migration is asymmetric, alleles spend more time in some habitats than others, disfavoring those that persist longer in unfavorable environments while favoring those that remain in favorable ones. Because such asymmetry can result from both increases and decreases in migration rates, either change can promote or hinder local adaptation. Notably, this dual effect emerges purely from the interplay between asymmetric migration and antagonistic selection, without invoking additional demographic or evolutionary mechanisms often proposed to explain how increased migration can facilitate local adaptation (Kawecki and Holt, 2002; Garant et al., 2007; Hendry, 2017, Ch. 5).

In this note, we address three key questions in the context of a single-locus model where a diploid population is subdivided into two connected habitats. First, under what patterns of migration and selection can local adaptation of two alleles be maintained or lost? Second, when an allele is fixed, under what conditions can a new mutation establish a polymorphism by becoming locally adapted? Third, how does local adaptation arise through the recurrent invasion of mutations with weak effects on a continuous phenotype? To answer these, we derive exact analytical conditions for local adaptation using the well-established two-habitat migration model, which captures the interaction between gene flow and selection in shaping allele dynamics (Bulmer, 1972; Christiansen and Feldman, 1975; Nagylaki, 1992; Bürger, 2000; Lenormand, 2002; Leturque and Rousset, 2002; Nagylaki and Lou, 2008; Yeaman and Otto, 2011; Lascoux et al., 2016; Tomasini and Peischl, 2018).

## Two-habitat migration model

The two-habitat migration model considers a single diploid locus in a population divided into two connected habitats, A and B, with sizes *N*_A_ and *N*_B_ that vary with migration rates, while the total population size *N* remains constant (*N*_A_ + *N*_B_ = *N*). Each generation includes viability selection within habitats, genotype-independent population regulation, reproduction, and migration.

The probabilities of genotype *g* surviving viability selection in habitats A and B are denoted by *V*_A,*g*_ and *V*_B,*g*_, respectively. Migration rates are represented as *m*_AB_ (the proportion of individuals in habitat A that migrated from habitat B) and *m*_BA_ (the proportion of individuals in habitat B that migrated from habitat A). The proportions of individuals remaining in their native habitats are given by *m*_AA_ = 1 *−m*_AB_ and *m*_BB_ = 1 *−m*_BA_. Habitats are completely isolated when *m*_AB_ = *m*_BA_ = 0 and fully mixed when *m*_AB_ = *m*_BA_ = 1*/*2, while setting one of the migration rates to 0 recovers continent-island models (Haldane, 1930; Wright, 1931; Nagylaki, 1992). We assume the population is at a migration-demographic equilibrium.

Whenever two alleles, *x* and *y*, segregate in the population, the population genetics of this system is fully described by the following recursion

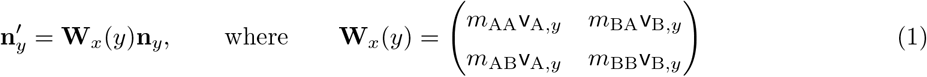

(Supporting information A). Here, **n**_*y*_ = (*N*_A,*y*_, *N*_B,*y*_) and 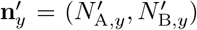 represents the local population sizes of *y* in consecutive generations in habitats A and B. The terms 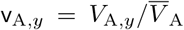 and 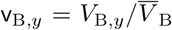 denote the relative probabilities of *y* surviving viability selection in habitats A and B, respectively. The survival probabilities of *y* in these habitats are given by *V*_A,*y*_ = *V*_A,*xy*_*p*_*x*|A_ + *V*_A,*yy*_*p*_*y*|A_ and *V*_B,*y*_ = *V*_B,*xy*_ *p*_*x*|B_ + *V*_B,*yy*_ *p*_*y*|B_, while 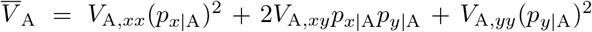 and 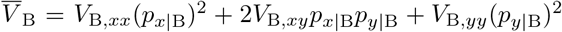 are the mean survival probabilities. The allele frequencies of *y* and *x* in habitat A are *p*_*y*|A_ and *p*_*x*|A_ = 1 *− p*_*y*|A_, while in habitat B, they are *p*_*y*|B_ and *p*_*x*|B_ = 1 *−p*_*y*|B_. The matrix **W**_*x*_(*y*) represents the fitness matrix of individuals carrying *y*, with each element quantifying the habitat-specific fitness of *y*. For instance, *m*_BA_v_B,*y*_ represents the expected number of offspring produced in A by a single *y* individual originating from B. We note that the model (eq. 1) is more commonly presented in terms of frequencies (e.g. Bulmer, 1972; Christiansen and Feldman, 1975; Nagylaki and Lou, 2008, see Supplementary information eq. A.14), but we use the above formulation as it more easily facilitates calculations of invasion fitness and weak selection approximations (see below and Supplementary information B).

## Local adaptation of a pair of alleles

Two alleles, *x* and *y*, are considered locally adapted when: (a) they experience antagonistic selection across habitats, such that, say, allele *x* is favored in habitat A (*V*_A,*yy*_ *< V*_A,*xy*_ *< V*_A,*xx*_) and *y* is favored in habitat B (*V*_B,*xx*_ *< V*_B,*xy*_ *< V*_B,*yy*_); and (b) they form a protected polymorphism (e.g., Kawecki, 2000; Kawecki and Holt, 2002). Protected polymorphisms occur when both alleles increase in frequency when rare, ensuring stable coexistence and preventing deterministic extinction due to frequency fluctuations. To investigate the conditions for local adaptation, we use invasion fitness functions, defined as the asymptotic per capita growth rate of a rare allele (Tuljapurkar, 1989; Metz et al., 1992). For the two-habitat migration model (eq. 1), the invasion fitness of *y* in a population of *x* is expressed as

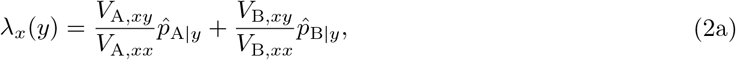

where 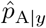 and 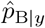 denote the probabilities of *y* being found in habitats A and B when rare. Their analytical expressions, which depend on migration rates and viability selection probabilities, are provided in Supporting Information B (eq. B.17). The influence of migration rates on 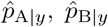, and thereby invasion fitness, is discussed in Section General analysis. Similarly, the invasion fitness of *x* in a population of *y* is

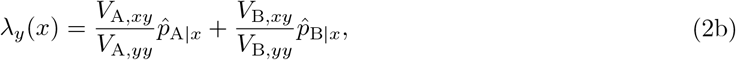

where 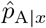 and 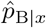 denote the probabilities of *x* being found in habitats A and B when it is rare. Whenever the invasion fitness of an allele is greater than 1, this allele can invade into the population, if smaller than 1, it can not. Whenever invasion fitness of both alleles exceed 1, the two alleles form a protected polymorphism. Therefore, under the assumption of antagonistic selection, *λ*_*x*_(*y*) *>* 1 and *λ*_*y*_(*x*) *>* 1 define the conditions for local adaptation of *x* and *y*. These conditions will be utilized in the general analysis (Section General analysis) and will be slightly refined for the second model (Section The emergence of local adaptation under weak selection).

## General analysis

Without loss of generality, we will simplify the notation in the above equations by setting *V*_A,*xy*_ = 1 = *V*_B,*xy*_, and *s*_A_ = *V*_A,*xx*_ *− V*_A,*xy*_ *>* 0, *t*_A_ = *V*_A,*xy*_ *− V*_A,*yy*_ *>* 0 and *s*_B_ = *V*_B,*xy*_ *− V*_B,*xx*_ *>* 0, *t*_B_ = *V*_B,*yy*_ *− V*_B,*xy*_ *>* 0, where *s*_A_, *t*_A_ quantify the selective advantages of *x* in habitat A and *s*_B_, *t*_B_ of *y* in habitat B, respectively. The conditions for local adaptation of alleles *x* and *y* (i.e., *λ*_*x*_(*y*) *>* 1 and *λ*_*y*_(*x*) *>* 1) can be expressed as

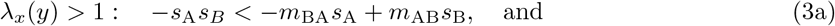

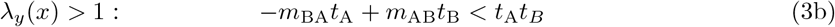

(Supporting information C). The first inequality (eq. 3a) specifies the condition under which *y* can adapt to habitat A (i.e., invade when rare), while the second inequality (eq. 3b) specifies when *x* can adapt to habitat B (i.e., invade when rare). If both inequalities are satisfied, the alleles *x* and *y* are locally adapted. Conversely, if either inequality is not satisfied, the corresponding allele cannot locally adapt. These conditions are illustrated in Figure 1.

**Figure 1:**
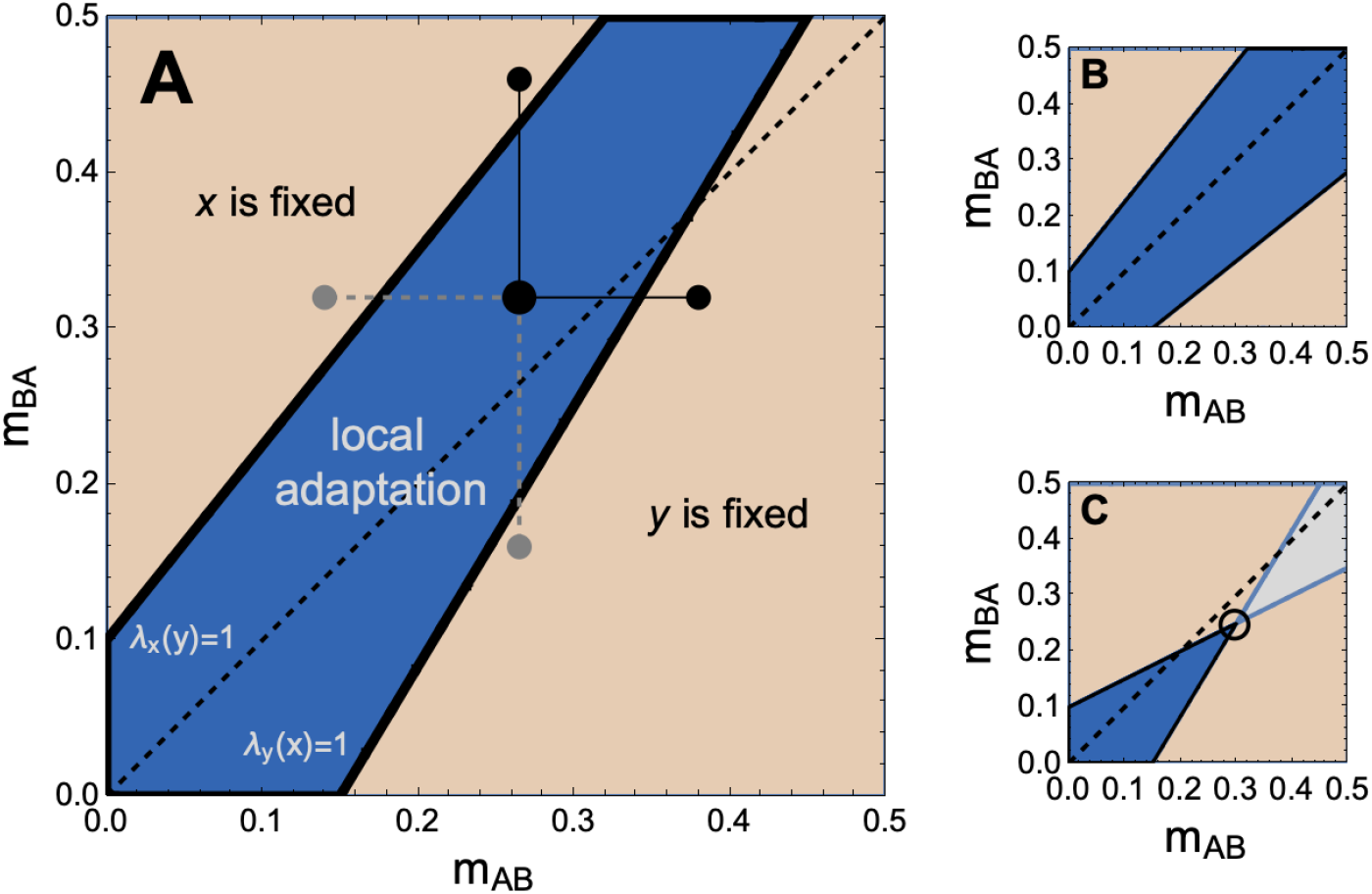
The conditions for local adaptation of two alleles, *x* and *y*, (eq. 3) are shown as a function of *m*_AB_ and *m*_BA_. In all panels, *s*_B_ = 0.1 and *t*_A_ = 0.15, while *s*_A_ and *t*_B_ vary between panels. The top thick black line represents the invasion boundary *λ*_*x*_(*y*) = 1, given by *m*_BA_ = *s*_B_ + *m*_AB_*s*_B_*/s*_A_, above which *y* cannot invade, leading to fixation of allele *x*. The bottom thick black line represents the invasion boundary *λ*_*y*_(*x*) = 1, given by *m*_BA_ = − *t*_B_ + *m*_AB_*t*_B_*/t*_A_, below which *x* cannot invade, leading to fixation of allele *y*. The blue region indicates where *x* and *y* form a protected polymorphism, defined by *λ*_*x*_(*y*) *>* 1 and *λ*_*y*_(*x*) *>* 1 (both inequalities in eq. 3 are satisfied). The black dashed line denotes symmetric migration rates (*m*_AB_ = *m*_BA_). **A)** Parameter values are *s*_A_ = 0.08 and *t*_B_ = 0.25. The black lines connecting black dots illustrate that increasing migration rates can lead to a loss of local adaptation. The gray dashed lines show that local adaptation can also be lost by decreasing migration rates. **B)** Parameter values are *s*_A_ = 0.08 and *t*_B_ = 0.12. In contrast to panel A, where increasing symmetric migration rates lead to the loss of local adaptation of *x*, local adaptation is maintained in this panel. **C)** Parameter values are *s*_A_ = 0.2 and *t*_B_ = 0.25. This panel demonstrates that invasion boundaries can intersect (empty circle), resulting in unprotected polymorphism of *x* and *y* near the intersection. The gray area indicates where *λ*_*x*_(*y*) *<* 1 and *λ*_*y*_(*x*) *<* 1.

We now address the first question posed in the introduction regarding the effect of migration on two locally adapted alleles, *x* and *y*. From eq. (3), it is evident that local adaptation can be lost as *m*_BA_ increases (Figure 1A, moving vertically from the large black dot to the small black dot), which selectively disfavors *y* (eq. 3a), or as *m*_AB_ increases (Figure 1A, moving horizontally from the large black dot to the small black dot), which selectively disfavors *x* (eq. 3b). Higher migration rates reduce the invasion fitness of these alleles by increasing the time they spend in their less favorable habitat, supporting the standard argument that gene flow hinders local adaptation. This occurs because changes in migration rates shift the balance of time alleles spend in each habitat. For example, increasing *m*_AB_ raises the proportion of individuals in A who are immigrants from B, thereby reducing the time (local) alleles remain in A and increasing the time they spend in B, where *x* is disfavored. This effect holds regardless of the strength of antagonistic selection. However, it is also evident that similar conclusions can be drawn by considering decreases in migration rates (Figure 1A, moving from the big black circle along gray dashed lines). Specifically, *y* is disfavored not only when *m*_BA_ increases but also when *m*_AB_ decreases (eq.3a), and *x* is disfavored not only when *m*_AB_ increases but also when *m*_BA_ decreases (eq.3b). In both cases, alleles spend more time in their disfavored habitats, which reduces their invasion fitness. Formally, these results can be understood by analyzing how the frequencies 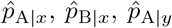, and 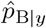 change with migration rates and how these changes impact the invasion fitnesses in eq. (2) (Supporting information B, eq. B.17). In our model, increasing *m*_AB_ always increases 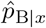 and 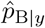, while increasing *m*_BA_ always increases 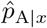 and 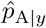; the opposite occurs when migration rates decrease.

The above discussion also addresses the second question posed in the introduction regarding the conditions under which a newly arising mutation can locally adapt. For example, in a population where *x* is fixed and the first inequality (eq. 3a) is violated, *y* cannot initially adapt (Figure 1A, gray dot in the left brown region). However, if *m*_AB_ increases (or *m*_BA_ decreases), the first inequality (eq. 3a) may eventually be satisfied, allowing *y* to adapt (Figure 1A, moving horizontally along the gray dashed line towards the large black dot). Further increases in *m*_AB_ (or decrease in *m*_BA_) could even lead to the loss of the initially fixed allele *x* if the inequality in eq. 3b is violated (Figure 1A, moving horizontally along the black line towards the small black dot). Increasing the migration rate *m*_AB_ (or decreasing *m*_BA_) can thus facilitate the local adaptation of an allele that is antagonistically favored in habitat B. The underlying logic is similar to the one above in that increasing *m*_AB_ raises the proportion of individuals in A who originate from B, which reduces the time alleles spend in A and extends their time in B, where *y* is favored. Similarly, in a population where *y* is fixed and the second inequality (eq. 3b) is violated, *x* cannot initially adapt Figure 1A, gray circle in the right brown region). However, as *m*_BA_ increases (or *m*_AB_ decreases), this favors the local adaptation of *x* and may eventually lead to the loss of the initially fixed allele *y* (Figure 1A, moving upwards from the gray circle towards the small black dot).

Both questions considered above demonstrate that migration rates play a dual role in influencing the balance between local adaptation and maladaptation: both increasing and decreasing migration rates can either facilitate or impede local adaptation. Yet this dual role does not apply if the two migration rates are symmetric. In such a case, an increase in migration causes both alleles to spend more time in their disfavoured habitats, thus reducing their invasion fitness and disfavouring local adaptation (Figure 1, dashed lines). Whether an allele is ultimately lost depends on the selection coefficients *s*_A_, *s*_B_, *t*_A_, and *t*_B_ (compare panels A and B, Figure 1). In the symmetric migration model, the condition for local adaptation of *y* (eq. 3a), as discussed by Lenormand (2002, p. 184), is recovered by setting *s*_A_ = *s* and *s*_B_ = *αs*, yielding *m/s < α/*(1 *− α*).

If the ratio *t*_B_*/t*_A_ is sufficiently larger than *s*_B_*/s*_A_, the two invasion boundaries defined by *λ*_*x*_(*y*) = 1 and *λ*_*y*_(*x*) = 1 intersect (Figure 1C, empty circle). This intersection implies that, in the vicinity of this point where either one or both alleles can not invade when rare (*λ*_*x*_(*y*) *<* 1 or *λ*_*y*_(*x*) *<* 1 or both), the two alleles can nevertheless form a stable polymorphism (Priklopil, 2012). However, because *λ*_*x*_(*y*) *<* 1 and/or *λ*_*y*_(*x*) *<* 1, large perturbations in allele frequencies can cause one of the alleles to go extinct. Thus, in the neighbourhood of the intersection, alleles *x* and *y* can form an *un*protected polymorphism, where the polymorphism persists but is not stable under large perturbations. We are unaware of any earlier results demonstrating unprotected polymorphism in the migration model (for special cases, see Novak, 2011; Priklopil, 2012). In this paper, we disregard such scenarios as instances of local adaptation, even though such polymorphisms may persist for significant periods of time.

## The emergence of local adaptation under weak selection

Local adaptation is generally not possible under weak selection. This can be seen from the inequalities in eq. (3): for *s*_A_ and *s*_B_ of order *ϵ* (where small *ϵ* implies weak selection), the left-hand side of eq. (3a), *−s*_A_*s*_B_, and the right-hand side of eq. (3b), *t*_A_*t*_B_, are both of order *ϵ*^2^, whereas the term *−m*_BA_*s*_A_+*m*_AB_*s*_B_ is of order *ϵ*. If this term is negative, the inequality in eq. (3a) fails; if positive, the inequality in eq. (3b) fails. Thus, under weak selection, both inequalities cannot generally hold simultaneously, and a single allele invariably fixes in the population (Supporting information C.2).

This result reflects a broader principle where any successfully invading mutation generally replaces the ancestral allele (classic results since Wright, 1931; Hamilton, 1964, see also Priklopil and Lehmann 2020, 2021). However, as mutations introduce new alleles, recurrent invasions and fixations follow the positive selection gradient (the slope of invasion fitness). This mutation-limited process eventually leads to the fixation of an allele near a so-called singular allele, where the selection gradient is zero. The nature of the singular allele determines the subsequent course of adaptive evolutionary dynamics. Because we are focused on the emergence of two locally adapted alleles, we require that the singular allele be an evolutionary branching point (EBP) (Geritz et al., 1998), a criterion previously used to assess local adaptation (e.g., Svardal et al., 2015). An EBP enables the formation of a protected polymorphism near the singular allele, satisfying *λ*_*x*_(*y*) *>* 1 and *λ*_*y*_(*x*) *>* 1, while resisting invasions by new mutations to prevent allele displacement and ensure long-term coexistence. In contrast, a continuously stable strategy (CSS) represents an evolutionary endpoint where such coexistence is not possible.

To analyze the emergence of local adaptation under a weak selection regime, we assume a continuum of potential alleles that contribute additively to a continuous phenotype. The population is assumed to be initially monomorphic with new alleles arising infrequently through mutations of small phenotypic effect leading to weak selection. The additive phenotype determines the survival probability of each genotype in its habitat. Specifically, the phenotypic contribution of an allele *x* is denoted by *ϕ*_*x*_, and when a mutation produces a new allele *y*, the phenotype of, for example, the heterozygote *xy* is given by *ϕ*_*xy*_ = (*ϕ*_*x*_ + *ϕ*_*y*_)*/*2. Homozygote phenotypes as well as phenotypes of any newly arising alleles are defined analogously. The dynamics of two alleles are governed by the recursion equations (eq. 1), with survival probabilities for genotype *g* defined as

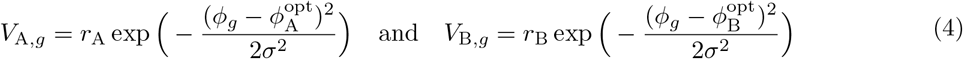

in habitats A and B, respectively, and where *r*_A_ and *r*_B_ are the maximum surviving probabilities in the respective habitats and *σ* controls the strength of viability selection. The phenotypic values 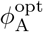 and 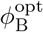 maximize the survival of individuals in habitats A and B, respectively, and by imposing that all individual phenotypes take values in between 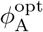 and 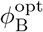 (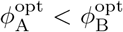), the selection is antagonistic. A special case of the model where *m*_AB_ + *m*_BA_ = 1 was analysed in Kisdi and Geritz (1999).

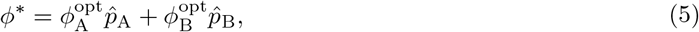

We find that singular allele gives rise to a singular phenotype where 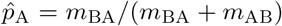 and 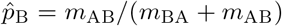 are the asymptotic probabilities to find a selectively neutral allele in habitats A and B, respectively, and can also be interpreted as the relative population sizes of the habitats (Supporting information C.2, eq. C.43). The singular phenotype is thus a weighted average of the optimal phenotypes in each habitat, with weights as relative habitat sizes. In the two-habitat migration model (eqs. 1 and 4), adaptive dynamics will always approach the singularity in eq. (5), which is an EBP whenever

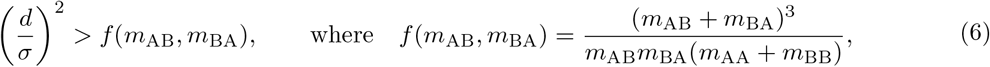

and where 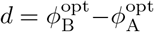 is the difference in habitat-specific optimal phenotypes. Whenever the condition in eq. (6) is satisfied, the adaptive dynamics of recurrent invasions and fixations of mutations with weak phenotypic effects will lead to local adaptation of two alleles. If the condition is not satisfied, the singular point is a CSS and long-term polymorphism is not possible (even though short-term polymorphisms are always possible in this model, EBP is not). We note that for the special case where *m*_AB_ + *m*_BA_ = 1, eqs. (5)-(6) simplify to the expressions given in Kisdi and Geritz (1999, eqs. 7-8).

The above results show that local adaptation depends on two key factors in eq. (6): the intensity of antagonistic selection *d/σ* (left-hand side) and the migration dynamics *f* (*m*_AB_, *m*_BA_) (right-hand side). Increasing the left-hand side favors local adaptation by strengthening antagonistic selection, which occurs when the distance between optimal phenotypes increases (*d*) or when viability probabilities narrow (*σ* decreases). The region of local adaptation (blue regions, Figure 2) shrinks with lower *d/σ* (compare panels A and B, Figure 2) and expands with higher *d/σ* (compare panels A and C, Figure 2).

**Figure 2:**
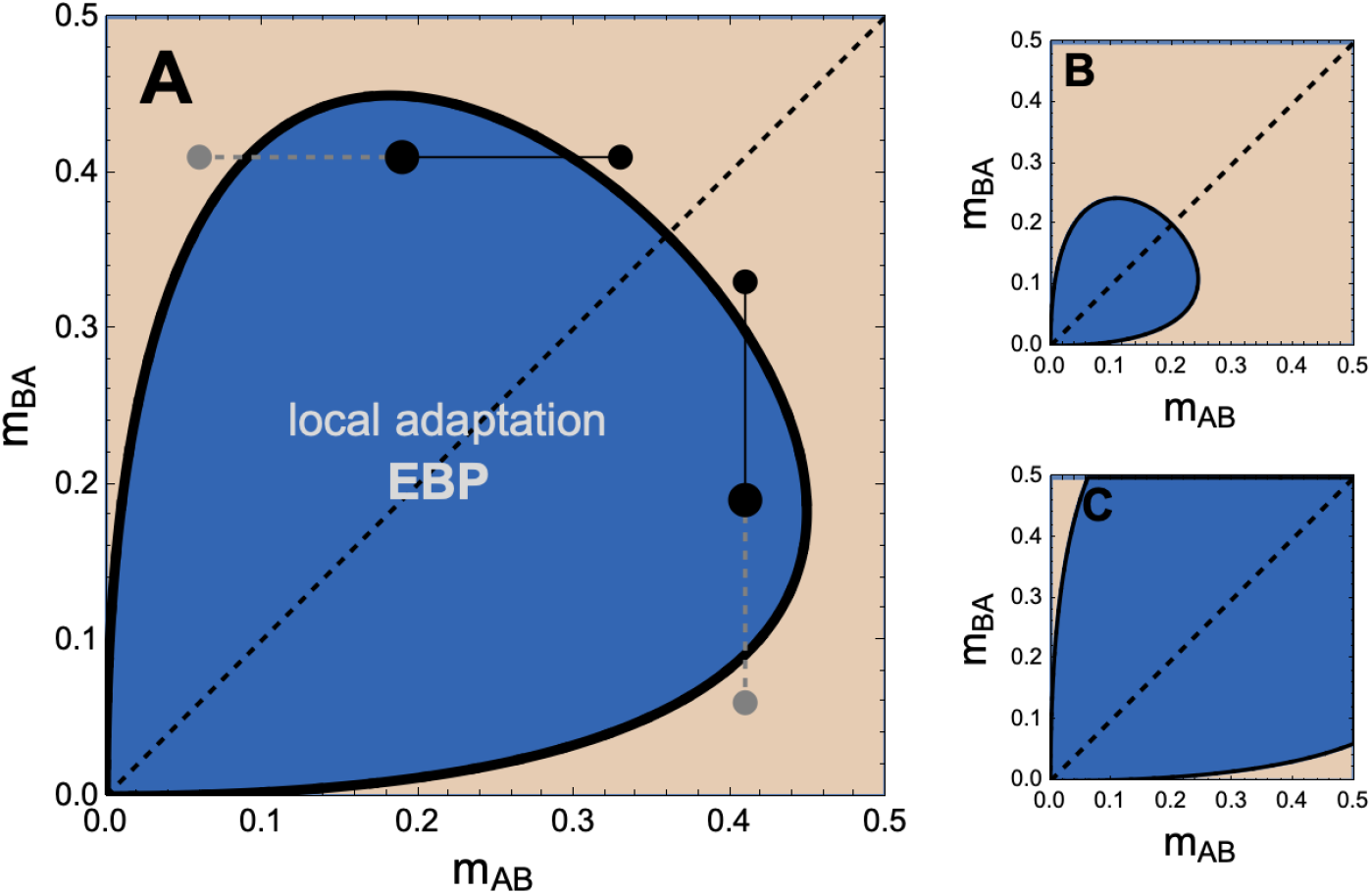
The condition for local adaptation, resulting from a gradual adaptive process of a continuous phenotype, as determined by eq. (6). Blue regions indicate (*m*_AB_, *m*_BA_) values where the singularity is an EBP (allowing local adaptation), while brown regions represents a CSS (evolutionary endpoint). The diagonal black dashed line denotes symmetric migration rates (*m*_AB_ = *m*_BA_). **A)** For *d/σ* = 1.5, an EBP transitions to a CSS as migration rates increase (moving from large black dots along black lines to small black dots) or decrease (moving from large black dots along gray dashed lines to small gray dots). **B)** For *d/σ* = 1, weaker antagonistic selection compared to panel A reduces the area supporting local adaptation. **C)** For *d/σ* = 2, stronger antagonistic selection compared to panel A increases the area supporting local adaptation. Unlike panels A and B, increasing symmetric migration rates does not result in the loss of local adaptation.

Reducing the right-hand side, *f* (*m*_AB_, *m*_BA_), also favors local adaptation. However, the function *f* (*m*_AB_, *m*_BA_) behaves non-monotonically with migration rates. Specifically, it increases with *m*_AB_ when 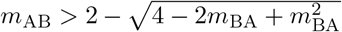, and decreases otherwise. Similarly, it increases with *m*_BA_ when 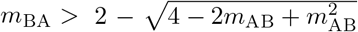, and decreases otherwise. This duality highlights that both increasing and decreasing migration rates can either favor or hinder local adaptation. For example, increasing *m*_AB_ can cause the loss of local adaptation (Figure 2A, moving from the large black dot horizontally to the right small black dot) because alleles spend more time in disfavored habitats, reducing invasion fitness. Conversely, decreasing *m*_AB_ can also lead to a loss of local adaptation (Figure 2A, moving from the large black dot horizontally to the left small gray dot) because alleles become largely confined to one habitat, favoring the fixation of a single allele. Similar arguments apply to increasing and decreasing *m*_BA_ (vertical transitions, Figure 2A). These conclusions are consistent with the general analysis presented above (Section General analysis).

Under symmetric migration, the above conclusions no longer hold, in agreement with the insights from the general analysis on symmetric migration. Under symmetric migration (*m*_BA_ = *m*_AB_ = *m*), the singular allele produces a singular phenotype located exactly halfway between the habitat-specific optimal phenotypes, 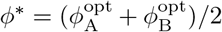 (eq. 5). This midpoint arises because migration is symmetric, causing both alleles to spend equal time in their favoured/disfavoured habitats. The condition for local adaptation under symmetric migration is given in eq. (6) with the right-hand side simplifying to 4*m/*(1 *− m*). Because this expression increases with migration rate *m*, it makes local adaptation progressively more difficult as *m* increases. Whether for large *m* local adaptation is possible or not depends on the parameter *d/σ* (compare panels A and B with panel C, Figure 2).

## Discussion

Migration plays a dual role in shaping local adaptation by either opposing or facilitating genetic differentiation. Through two models – one analyzing arbitrary antagonistic selection and the other examining gradual adaptation in a continuous phenotype – we demonstrate that asymmetric migration alone, by altering the time alleles spend in each habitat, can explain these outcomes without invoking additional mechanisms.

Under symmetric migration, increasing migration reduces local adaptation as alleles spend more time in less favorable non-local habitats, lowering their mean fitness. In contrast, asymmetric migration produces dual effects. Increasing *m*_AB_ raises the proportion of migrants in A, which decreases the time local alleles remain in A. As a result, alleles spend more time in B, favoring the allele that is advantageous there while disfavoring the other. The opposite effect occurs when *m*_BA_ increases. Reducing migration rates produces similar effects. These findings challenge the conventional view that increasing migration generally opposes local adaptation, instead highlighting the critical role of asymmetry. The dual role of migration rates described above is likely to extend to other models, as long as changes in migration rates lead to monotonic shifts in where alleles spend their time.

Our analysis makes several additional contributions. While the conditions for protected polymorphisms in two-habitat migration models have been established since Bulmer (1972), we identify novel cases of unprotected polymorphisms. In such cases, a pair of alleles can stably coexist, but since at least one allele cannot increase in frequency when rare, the polymorphism is not protected from large fluctuations in allele frequencies and is unlikely to evolve through the invasion of new rare mutations. Additionally, we extend the adaptive dynamics framework to this model, addressing challenges in analyzing second-order effects in structured populations. Using eigenvector perturbation methods (see Supporting Information B), we provide a complete analytical treatment and show that the Levene model analyzed by Kisdi and Geritz (1999) is a special case of our results. Finally, unlike previous studies, our analysis focuses on forward-in-time dynamic equations rather than backward-in-time processes based on reproductive values (e.g., Leturque and Rousset, 2002).

This work offers a comprehensive understanding of the dual role of migration in shaping local adaptation, providing novel insights into the interplay between migration asymmetry, selection, and population structure. By identifying scenarios where migration can simultaneously promote and hinder adaptation and by uncovering unprotected polymorphisms in two-habitat models, this study advances theoretical frameworks critical for understanding genetic differentiation in heterogeneous environments.

## Supporting information

### A. Two-habitat migration model

Here, we derive the migration model from the main text, previously analyzed in Smith, 1970; Bulmer, 1972; Christiansen and Feldman, 1975; Bürger, 2000; Nagylaki and Lou, 2008. The model considers a population subdivided into two connected habitats, A and B, with allele population sizes *N*_A_ and *N*_B_, respectively, which can change due to migration while the total population size remains fixed (*N*_A_ +*N*_B_ = *N*). We assume non-overlapping generations and a single diploid locus.

At the start of each generation, individuals undergo habitat-specific viability selection. The survival probability of a genotype *g* is *V*_A,*g*_ in habitat A and *V*_B,*g*_ in habitat B. When two alleles *x* and *y* segregate, the survival probabilities of, for example, allele *y* are

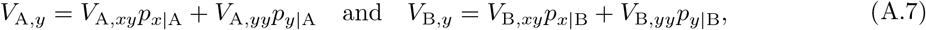

in habitats A and B, respectively, where *p*_*y*|A_ and *p*_*x*|A_ = 1 *− p*_*y*|A_ are the frequencies of *y* and *x* in habitat A, and similarly, *p*_*y*|B_ and *p*_*x*|B_ = 1 *− p*_*y*|B_ in habitat B. The average survival probabilities are 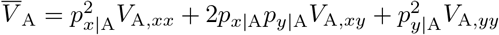and 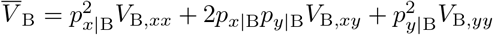.

After viability selection, all survivors become reproducing adults. They mate randomly and produce a large number of offspring, which undergo non-selective competition to restore the habitat-specific population sizes. Habitat sizes thus satisfy 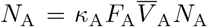 and 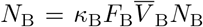, where *F*_A_ and *F*_B_ are habitat-specific fecundities, and *κ*_A_ and *κ*_B_ describe the non-selective competition within each habitat.

Following reproduction and population regulation, offspring either remain in their native habitat or disperse to the alternate habitat. The probability of dispersal is independent of genotype, and the forward migration rate, for example, from B to A is denoted with 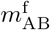, with 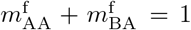 and 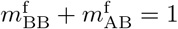. Backward migration rates, which depend on local population sizes, are defined as

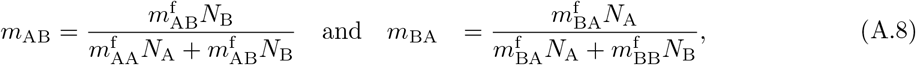

where *m*_AB_ (or *m*_BA_) is the probability that a randomly sampled individual in A (or B) originates from B (or A). Conservation of individuals ensures *m*_AA_ + *m*_AB_ = 1 and *m*_BA_ + *m*_BB_ = 1. After dispersal, the life cycle restarts.

In the following sections, we first describe the demographic process of migration (Section A.1), followed by the population genetics of two alleles under demographic equilibrium (Section A.2).

#### A.1. The demographic process of migration

Here, we consider the dynamics of the demographic process of migration and its steady state. Because viability selection, reproduction, and population regulation preserve the local population sizes, and migration of an individual is assumed to be independent of its genotype, the local (allele) population sizes **n** = (*N*_A_, *N*_B_) follow the recursion

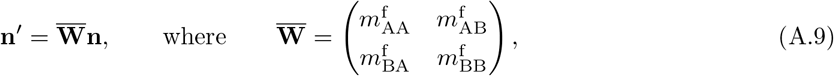

and where 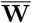 is the (average) fitness matrix of an (average) allele in the population. We refer to this system as a neutralized system because alleles following these dynamics behave like average alleles (averaged over all segregating alleles in the population), effectively acting as if they were neutral. In the two-habitat migration model, the neutralized and neutral systems (where all alleles have identical survival probabilities) are equivalent, though this is not generally the case (Priklopil and Lehmann, 2024). Because the total population is fixed, we have 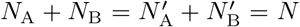. Since the migration matrix is stochastic, **N** converges to a unique line of equilibria 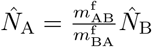, and we have

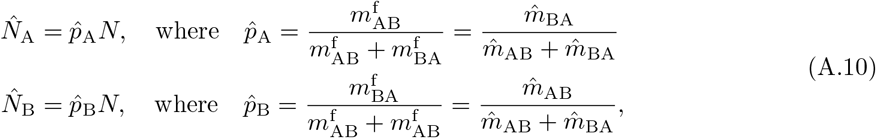

where 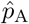 and 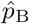 represent the relative habitat sizes of A and B, respectively, at demographic steady state. Their expressions follow from the identity 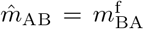 and 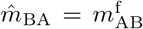, which holds at the demographic steady state and can be derived using eq. (A.8).

We note that eq. (A.9) describes a forward-in-time demographic process, where 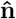 is the right eigenvector of 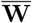 associated with the dominant eigenvalue 1 (the corresponding left eigenvector is *k*(1, 1) for any constant *k*). The backward-in-time version of this process, as discussed e.g. in Leturque and Rousset (2002) and in Rousset (2004, pg., 153), uses the left and right eigenvectors of the forward-in-time process in reverse roles. While backward migration rates depend on local population sizes (denoted at demographic steady state with a hat-notation), forward migration rates are individual-level characteristics and independent of population sizes. Throughout the main text and the Supporting information, we assume that the population is always at demographic steady state (eq. A.10), as the demographic process is independent of genotypic variation and stabilizes before genetic variation arises. Consequently, we omit the hat notation and use the simplifications 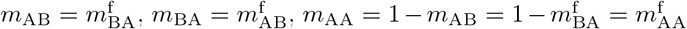 and 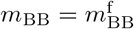.

At demographic steady state (eq. A.10), the local population sizes *N*_A_ and *N*_B_ are equal only if migration rates are symmetric 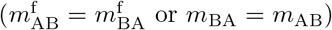; otherwise, they differ. The demographic equilibrium can also be expressed as 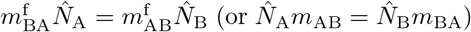, indicating a balance in the number of migrants moving between habitats. This equality ensures conservation of migrant numbers across habitats, even if migration rates differ. However, the proportion of immigrants is generally unequal between habitats (*m*_AB ≠_ *m*_BA_). The total number of migrants from A to B in each generation is *N*_A_*m*_AB_ = *Nm*_BA_*m*_AB_*/*(*m*_AB_ + *m*_BA_), with the same number migrating from B to A. Consequently, the total proportion of migrating individuals (out of *N*) is

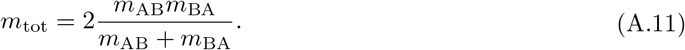

This proportion increases as either *m*_AB_ or *m*_BA_ increases, indicating that higher migration rates lead to more total migrants. For symmetric migration (*m*_AB_ = *m*_BA_ = *m*), the total migrating proportion simplifies to *m*. When *m*_AB_ = *m*_BA_ = 1*/*2, half of the population migrates each generation (*m*_tot_ = 1*/*2). Conversely, when *m*_AB_ = 0 = *m*_BA_, no migration occurs, and the population remains in its initial state.

### A.2. Population genetics of two alleles

Here, we present the population genetics of two segregating alleles *x* and *y* in a population that is at a demographic steady state (eq. A.10, Section A.1). Because local population sizes are fixed at this steady state at all times, it is sufficient to analyse the dynamics of only one of the two alleles, say the mutant allele *y*. Given the life-cycle summarized above, the population genetics can thus be fully expressed with a single recursion

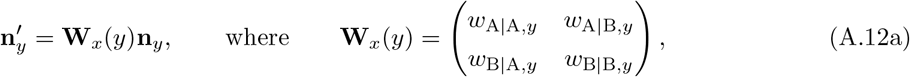

where **n**_*y*_ = (*N*_A,*y*_, *N*_B,*y*_) are the local population sizes of *y* across the two habitats, and **W**_*x*_(*y*) is the (individual) fitness matrix of (an allele) *y*. Each element in the fitness matrix gives the class-specific fitness of *y*, for example, *w*_A|B,*y*_ is defined as the expected number of offspring produced into A by a single *y* in B. The class-specific fitnesses can be expressed as

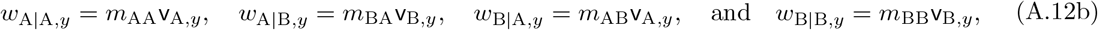

where

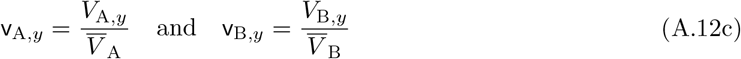

are the relative survival probabilities of allele *y* in habitats A and B, respectively. Note that, for example, the interpretation of

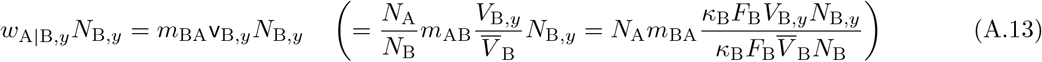

where we used eq. (A.10), is the number of individuals in A in the offspring generation (*N*_A_) that came from B in the parent generation (*N*_A_*m*_BA_), of which the fraction 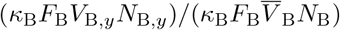 are *y* after viability selection, reproduction and population regulation. This thus gives the total number of *y* in A that came from B in the previous generation, hence justifying *w*_A|B,*y*_ as the class-specific individual fitness.

The population genetics of two alleles (eq. A.12b) has been more commonly expressed in terms of allele frequencies as

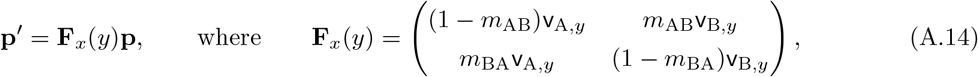

where **p** = (*p*_*y*|A_, *p*_*y*|B_) is a vector of habitat-specific frequencies of *y* (see e.g., Leturque and Rousset, 2002). We note that eq. (A.14) defines the backward-in-time version of the forward-in-time process described in eq. (A.12b), and thus **F**_*x*_(*y*) and **W**_*x*_(*y*) are adjoint matrices of each other. While all results can be derived using either matrix, in this paper, we derive our results from the forward-in-time recursion (eq. A.12b), as it allows for a more unified presentation of the strong and weak selection results under a single analysis, *sensu* Priklopil and Lehmann (2024).

## B. Invasion fitness

Invasion fitness of an allele is its asymptotic growth rate when rare, which in our model is equal to the dominant eigenvalue of an individual fitness matrix linearized around an equilibrium where this allele is absent. The invasion fitness *λ*_*x*_(*y*) of *y* in a population of *x* satisfies

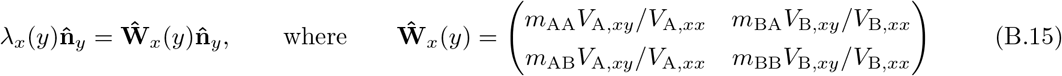

is the invasion fitness matrix, calculated by linearizing the fitness matrix (eq. A.12) about the trivial equilibrium where *y* is absent, and 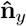 is the right eigenvector associated with *λ*_*x*_(*y*). By premultiplying the left-hand side by an arbitrary vector, one can obtain an explicit expression for invasion fitness. Here, we follow Priklopil and Lehmann (2024) and premultiply by the individual reproductive values 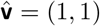 of a neutralized system (eq. A.9), because the resulting product gives us a measure of selection also in non-asymptotic dynamics and simplifies the weak selection approximation in Section B.1. The invasion fitness then simplifies to

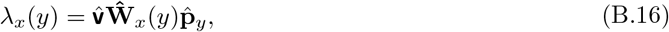

where 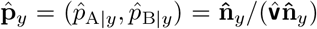 is a vector of probabilities to find the rare allele *y*, asymptotically, in habitats A and B. It can be re-written as

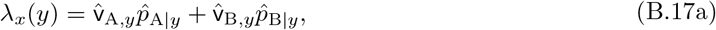

where 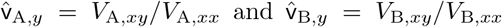, and where the elements of the dominant right eigenvector are

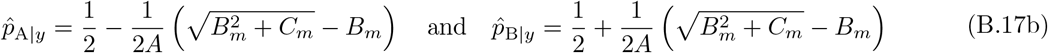

with 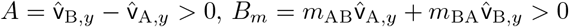, and

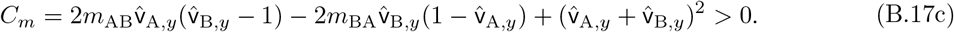

Note that if it were not for the term *C*_*m*_, the term in the brackets would be 0. Therefore, and because *A* is independent of the migration rate, increasing or decreasing *C*_*m*_ by increasing or decreasing *m*_AB_ or *m*_BA_ fully determines whether the element of the right eigenvector increases or decreases. Specifically, increasing *m*_AB_ increases *C*_*m*_, and hence 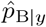 increases and 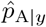 decreases, whereas the opposite occurs by increasing *m*_BA_.

Using an analoguous invasion matrix for allele *x*, the invasion fitness of an allele *x* in a population of *y* can be expressed as

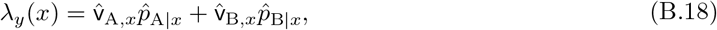

where 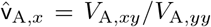 and 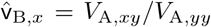, and where 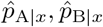 are the elements of the right eigenvector of the invasion matrix of a rare *x* corresponding to the dominant eigenvalue.

### B.1. Approximation under weak selective effects on a continuous phenotype

Let us now suppose that selection acts on a continuous phenotype *ϕ*, to which alleles contribute additively with small phenotypic effects. That is, we denote with *ϕ*_*g*_ the phenotype of a genotype *g* so that, for example, the phenotype of a heterozygote *xy* is *ϕ*_*xy*_ = (*ϕ*_*x*_ + *ϕ*_*y*_) where *ϕ*_*x*_ and *ϕ*_*y*_ are the phenotypic contributions of alleles *x* and *y*. We denote with *δ* = *ϕ*_*y*_ *− ϕ*_*x*_ the difference between the phenotypic contributions, and small *δ* reflects small mutational effect. Whenever the context is clear, we use *δ* instead of *y, x* in the arguments of functions. For example, we write 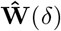 instead of 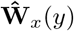 and *λ*(*δ*) instead of *λ*_*x*_(*y*), but noting that *δ* = 0 implies the expression still depends on *x*. The viability fitness functions for genotype *g* are given in the main text (eq. 4), but we replicate them here for completeness,

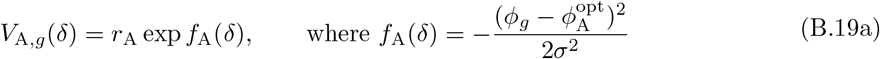

and

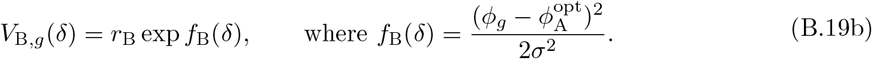

When the effect of mutations is small (*δ* is small), the invasion fitness of *y* in a resident population *x* (eq. B.16 or B.17) can be approximated up to second order as

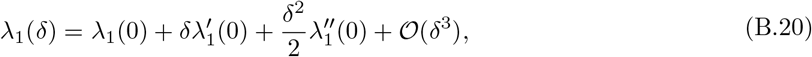

where

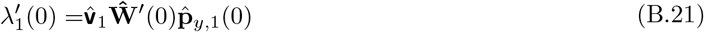

and

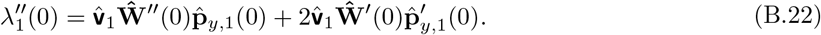

Here,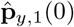 is the dominant right eigenvector of 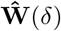 evaluated at *δ* = 0 and normalized such that 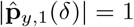 for all *δ*, and 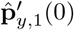 is its derivative at *δ* = 0. The vector 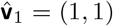 is the left dominant eigenvector of the neutralized system (eq. A.9) and is independent of *δ* (see above). We note that in this section, but nowhere else in the text, we enumerate the eigenvectors and eigenvalues due to the use of dominant as well as subdominant eigenvectors and eigenvalues.

Next, using only the invasion matrix (eq. B.15) and its derivatives evaluated at *δ* = 0, we will calculate all the eigenvectors and their perturbations needed for the weak selection approximation of invasion fitness up to second order (eqs. B.20-B.22). The expressions that are needed are the invasion matrix evaluated at *δ* = 0,

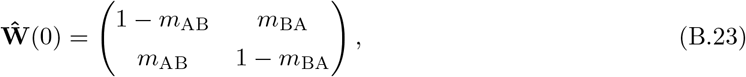

as well as its first-order derivative at *δ* = 0,

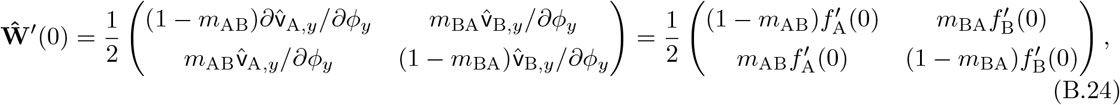

and the second-order derivative at *δ* = 0,

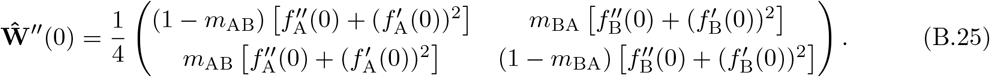

The factors 1*/*2 and 1*/*4 in front of the matrices arise from accounting for diploid individuals (e.g.,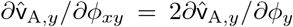. We also note that the neutral, or resident, fitness matrix (eq. B.23) is identical to the neutralised matrix (eq. A.9), and their eigenvalues are

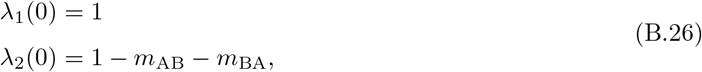

where *−*1 *< λ*_2_(0) *<* 1 is the subdominant eigenvalue. The left and right eigenvectors associated to *λ*_1_(0) are

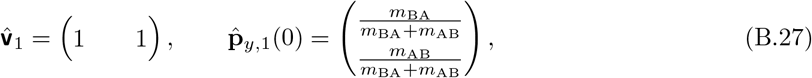

respectively. As a final step, we calculate the perturbation 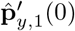 needed for eq. (B.22). This can be obtained, for example, by following the procedure given in Wilkinson, 1965, p. 69, Stewart 2001, Section 3.2, eq. 3.14; see also Caswell 2000, eq. 9.131. In this procedure we express the perturbation in terms of subdominant eigenvalues and eigenvectors as

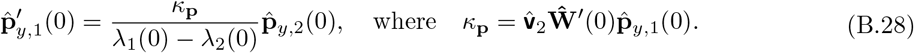

The right and left eigenvectors of the resident or neutralised matrix (eq. B.23 or eq. A.9) associated to the subdominant eigenvalue *λ*_2_(0) are

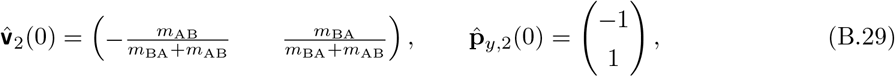

normalized such that their dot product is 1. The eigenvector-perturbation (eq. B.28) is thus given as

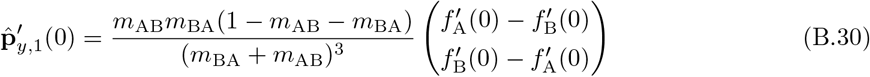

obtained by substituting eqs. (B.24), (B.27), and (B.29) into eq. (B.28).

We have now all the ingredients for the eigenvalue approximation (eqs. B.20-B.22). The expression for the first-order term is obtained by substituting eqs. (B.24) and (B.27) into eq. (B.21),

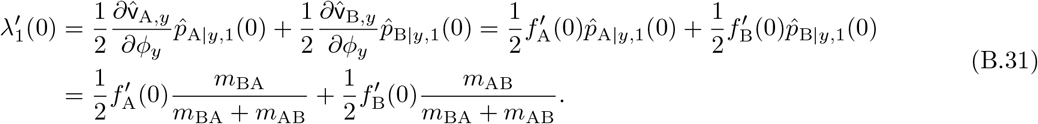

The expression for the second-order term is obtained by substituting eqs. (B.24)-(B.25), (B.27), and (B.30) into eq. (B.22),

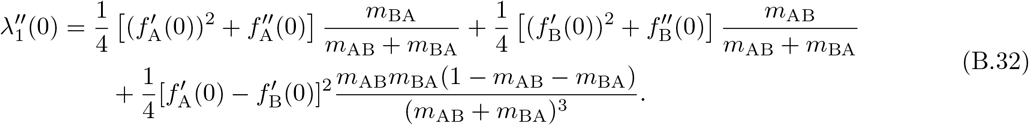

#### Symmetric migration

In the simpler case where migration is symmetric, *m*_BA_ = *m*_AB_ = *m*, we have 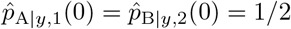, and the first-order condition (eq. B.31) becomes

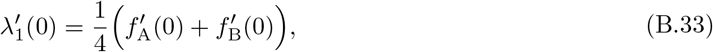

and the second-order condition (eq. B.32) becomes

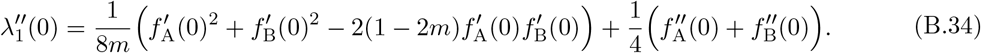

Note that whereas the second-order condition depends on *m*, the first-order condition is independent of migration.

## C. Conditions for local adaptation

Here we derive conditions for local adaptation of two alleles for the model with arbitrary patterns of viability selection (Section C.1) and for the model where mutations have a weak selective effect on a continuous phenotype (Section C.2).

### C.1. Arbitrary patterns of viability selection

Two alleles *x* and *y* are said to be locally adapted, if they coexist as a protected polymorphism (*λ*_*x*_(*y*) *>* 1 and *λ*_*y*_(*x*) *>* 1) under antagonistic selection. The expressions for invasion fitness’s are rather complex (eqs. B.17-B.18), however, and so we here derive a simple proxy for *λ*_*x*_(*y*) by following Bulmer (1972, proxy for *λ*_*y*_(*x*) can be derived similarly). First, we note that because all elements in the invasion fitness matrix (eq. B.15) are positive, the Perron-Frobenius theorem ensures the existence of a real and positive dominant eigenvalue that is greater than 0. Second, we note that all eigenvalues can be found by solving det 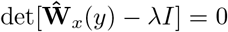, which leads to the characteristic equation *λ*^2^ *−* trace 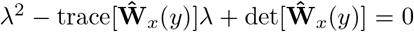 . This characteristic equation can be usefully re-written as

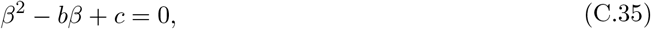

where *β* = *λ −* 1 and

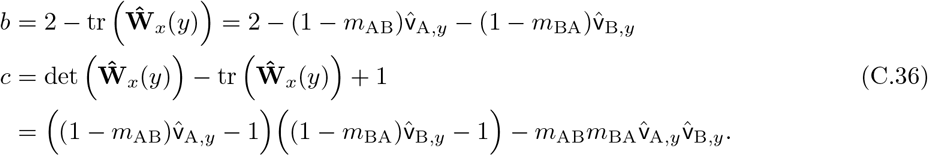

Since the quadratic term in eq. (C.35) has a positive coefficient (the coefficient is 1), the parabola in eq. (C.35) opens upward, and the coefficients *b* and *c* determine the signs of the roots (*β* = *λ −* 1) of eq. (C.35). Specifically, if *c <* 0, one root is positive and one is negative. If *c >* 0 and *b <* 0, then both roots are positive. If *c >* 0 and *b >* 0, both roots are negative. Thus, the dominant eigenvalue is greater than 1 whenever *c <* 0 or *b <* 0. However, we observe that if *c >* 0, then *b* is always positive. This occurs because for *c* to be positive, the first term on the right hand side on the final line in eq. (C.36) must be positive, and hence necessarily 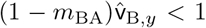, because by assumption of antagonistic selection we have 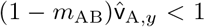. This in turn means that *b <* 0. Therefore, the sign of *c* alone determines whether the dominant eigenvalue is greater than 1. In other words, if *c >* 0, then *b* is always positive, and consequently, *λ*_*x*_(*y*) *<* 1. If *c <* 0, then *λ*_*x*_(*y*) *>* 1, regardless of the sign of *b*.

The condition for *c <* 0 in eq. (C.36) can be expressed as

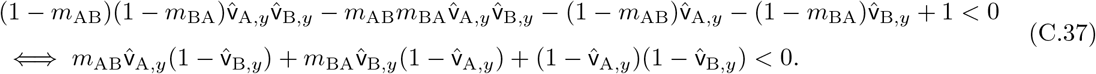

Dividing both sides of the final inequality in eq. (C.37) by 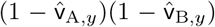 and reorganising, the condition *c <* 0 becomes

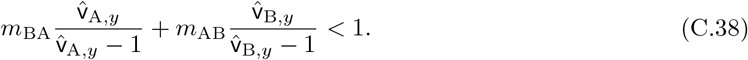

We have thus obtained that whenever eq. (C.38) is satisfied, then *λ*_*x*_(*y*) *>* 1 and otherwise *λ*_*x*_(*y*) *<* 1. Similar calculations hold to find a proxy for *λ*_*y*_(*x*), which is greater than 1 whenever

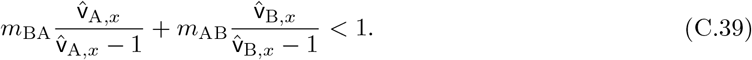

Condition for *x* and *y* to form a protected polymorphism is thus whenever both inequalities in eqs. (C.38)-(C.39) are satisfied. We note that these are exactly the conditions derived in Bulmer (1972, eq. 21) and in Nagylaki and Lou (2008, eq. 4.35) if we set *m*_AB_ = *m*_1_, *m*_BA_ = *m*_2_, *V*_A,*xy*_ = *V*_B,*xy*_ = 1, *V*_A,*xy*_ *−V*_A,*xx*_ = *s*_1_, *V*_A,*xy*_ *− V*_A,*yy*_ = *t*_1_, *V*_B,*xy*_ *− V*_B,*xx*_ = *s*_2_, *V*_B,*xy*_ *− V*_B,*yy*_ = *t*_2_ with *s*_1_ *<* 0 *< s*_2_ and *t*_1_ *<* 0 *< t*_2_. The condition eq. (C.38) is then equivalent to *m*_1_*/s*_1_ +*m*_2_*/s*_2_ *<* 1 and condition eq. (C.39) to *m*_1_*/t*_1_ +*m*_2_*/t*_2_ *<* 1.

#### Simplifying notation

Without loss of generality, we will simplify the notation in the above equations by setting *V*_A,*xy*_ = 1 = *V*_B,*xy*_, and *s*_A_ = *V*_A,*xx*_ *− V*_A,*xy*_ *>* 0, *t*_A_ = *V*_A,*xy*_ *− V*_A,*yy*_ *>* 0 and *s*_B_ = *V*_B,*xy*_ *− V*_B,*xx*_ *>* 0, *t*_B_ = *V*_B,*yy*_ *− V*_B,*xy*_ *>* 0, where *s*_A_, *t*_A_ quantify the selective advantages of *x* in habitat A and *s*_B_, *t*_B_ of *y* in habitat B, respectively. The condition for local adaptation of alleles *x* and *y* can then be stated as

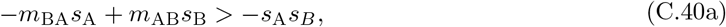

giving the condition for *y* being able to adapt to A (i.e., to invade into the population when rare, and hence is sign-equivalent with eq. C.38), and

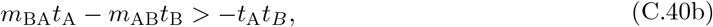

giving the condition for *x* being able to adapt to B (i.e., to invade into the population when rare, and hence is sign-equivalent with eq. C.39). By assuming that viability fitness functions are approximately linear in terms of the selection coefficients (which holds at least for small coefficients), so that *s*_A_ = *t*_A_ and *s*_B_ = *t*_B_, then the condition for protected polymorphism of *y, x* can be stated as

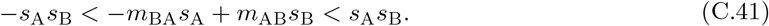

The first inequality is the invasion boundary of allele *y* and the second inequality the invasion boundary of *x*. This case has been referred to as the case of “no dominance” (Nagylaki and Lou, 2008, eq. 4.35).

### C.2. Weak selective effects on a continuous phenotype

The dominant eigenvalue *λ*_1_(*δ*) of the invasion fitness matrix (eq. B.15) determines whether a mutation can invade into the population. If the mutation can invade, and provided it is not eliminated in the initial phase of invasion due to demographic stochasticity, it will generically replace its ancestral resident allele and fix in the population (Rousset, 2004; Geritz, 2005; Dercole and Rinaldi, 2008; Priklopil and Lehmann, 2020). Recurrent mutation-invasion-fixation events then produce a step-by-step change in the phenotype until it reaches the vicinity of its *singular* value (the so-called candidate ESS), which can be found by solving eq. (B.31). In an alternative scenario, the phenotype reaches the boundary of its feasible phenotype space. However, in this paper, all phenotypes are assumed to be feasible, and we disregard such scenarios. Whether the adaptive process converges toward (or diverges away from) a singularity, is determined by the criterion for convergence stability (Eshel, 1983; Lessard, 1990, see below).

After the evolutionary process reaches the vicinity of a convergence stable phenotype *ϕ*^∗^, provided it exists, two qualitatively different evolutionary outcomes are possible. The first possibility is that *ϕ*^∗^ serves as the endpoint of the adaptive evolutionary process because selection near this singularity is stabilizing, and any emerging genetic variation will eventually be depleted in this mutation-limited evolutionary process (in our model, such “transient” polymorphisms are always possible as singularity is for all parameter values attached to the area of mutual invasion (Geritz et al., 1998). Calculations are not shown here). Such a singularity, where no long-term polymorphism is possible, is called a continuously stable ESS (CSS). The second possibility is that the convergence stable singularity *ϕ*^∗^ is an evolutionary branching point (EBP). In this case, selection near this singularity is disruptive, and any emerging genetic variation can persist indefinitely as protected genetic polymorphisms (Geritz et al., 1998).

We are therefore interested in finding conditions under which singularities are EBPs, as this provides the condition for local adaptation as the gradual evolutionary outcome (used e.g., in Svardal et al., 2015). EBPs can be identified algebraically by solving

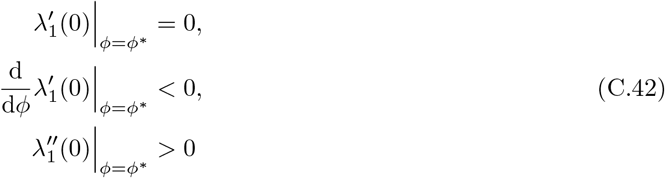

(Geritz et al., 1998; Della Rossa et al., 2015). If the final inequality has an opposite sign the singular *ϕ*^∗^ is a CSS.

Let us now apply the assumptions made on viability selection in eq. (4) (or eq. B.19). From the first equality in eq. (C.42) we find the singular phenotype by solving eq. (B.31),

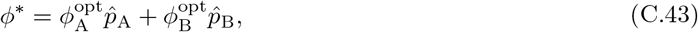

with 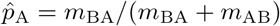 and 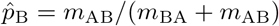 (see eqs. B.27 and A.10). This singularity is always convergence stable because the second line in eq. (C.42) gives us *−*1*/*(2*σ*^2^) *<* 0 which is always satisfied. Finally, using eq. (B.32), the singularity is an EBP (eq. C.42) whenever

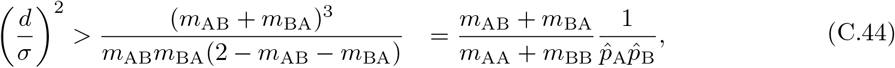

where 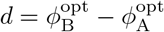. The right-hand side increases with *m*_AB_ whenever

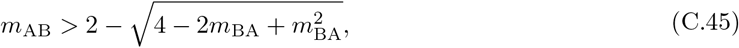

otherwise the right-hand side decreases. Similarly, the right-hand side increases with *m*_BA_ whenever

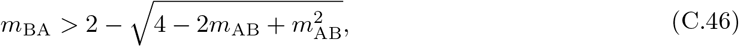

otherwise the right hand side decreases.

## References

Blanquart, F., Kaltz, O., Nuismer, S. L., and Gandon, S. (2013). A practical guide to measuring local adaptation. Ecology letters, 16(9):1195–1205.

Bulmer, M. (1972). Multiple niche polymorphism. The American Naturalist, 106(948):254–257.

Bürger, R. (2000). The Mathematical Theory of Selection, Recombination, and Mutation. John Wiley and Sons, New York.

Caswell, H. (2000). Matrix Population Models. Sinauer Associates, Massachusetts.

Christiansen, F. B. and Feldman, M. W. (1975). Subdivided populations: a review of the one-and two-locus deterministic theory. Theoretical Population Biology, 7(1):13–38.

Della Rossa, F., Dercole, F., and Landi, P. (2015). The branching bifurcation of adaptive dynamics. International Journal of Bifurcation and Chaos, 25(07):1540001.

Dercole, F. and Rinaldi, S. (2008). Analysis of Evolutionary Processes: The Adaptive Dynamics Approach and Its Applications. Princeton University Press, Princeton, NJ.

Eshel, I. (1983). Evolutionary and continuous stability. Journal ot Theoretical Biology, 103:99–111.

Garant, D., Forde, S. E., and Hendry, A. P. (2007). The multifarious effects of dispersal and gene flow on contemporary adaptation. Functional Ecology, pages 434–443.

Geritz, S. A. (2005). Resident-invader dynamics and the coexistence of similar strategies. Journal of mathematical biology, 50(1):67–82.

Geritz, S. A. H., Kisdi, E., Meszéna, G., and Metz, J. A. J. (1998). Evolutionarily singular strategies and the adaptive growth and branching of the evolutionary tree. Evolutionary Ecology, 12:35–57.

Haldane, J. (1930). A mathematical theory of natural and artificial selection.(part vi, isolation.). In Mathematical proceedings of the Cambridge philosophical society, volume 26, pages 220–230. Cambridge University Press.

Haldane, J. (1948). The theory of a cline. Journal of genetics, 48:277–284.

Hamilton, W. D. (1964). The genetical evolution of social behaviour, 1. Journal of Theoretical Biology, 7:1–16.

Hendry, A. P. (2017). Eco-evolutionary dynamics. Princeton university press.

Kawecki, T. J. (2000). Adaptation to marginal habitats: contrasting influence of the dispersal rate on the fate of alleles with small and large effects. Proceedings of the Royal Society of London. Series B: Biological Sciences, 267(1450):1315–1320.

Kawecki, T. J. and Ebert, D. (2004). Conceptual issues in local adaptation. Ecology letters, 7(12):1225– 1241.

Kawecki, T. J. and Holt, R. D. (2002). Evolutionary consequences of asymmetric dispersal rates. The American Naturalist, 160(3):333–347.

Kisdi, É. and Geritz, S. A. (1999). Adaptive dynamics in allele space: evolution of genetic polymorphism by small mutations in a heterogeneous environment. Evolution, 53(4):993–1008.

Lascoux, M., Glémin, S., and Savolainen, O. (2016). Local adaptation in plants. eLS, 25270:1–7.

Lenormand, T. (2002). Gene flow and the limits to natural selection. Trends in ecology & evolution, 17(4):183–189.

Lessard, S. (1990). Evolutionary stability: one concept, several meanings. Theoretical Population Biology, 37:159–170.

Leturque, H. and Rousset, F. (2002). Dispersal, kin competition, and the ideal free distribution in a spatially heterogeneous population. Theoretical Population Biology, 62:169–180.

Metz, J. A. J., Nisbet, R. M., and Geritz, S. A. H. (1992). How should we define fitness for general ecological scenarios? Trends in Ecology and Evolution, 7:198–202.

Nagylaki, T. (1992). Introduction to Population Genetics. Springer-Verlag, Heidelberg.

Nagylaki, T. and Lou, Y. (2008). The dynamics of migration–selection models. In Tutorials in mathematical biosciences IV: Evolution and ecology, pages 117–170. Springer.

Novak, S. (2011). The number of equilibria in the diallelic levene model with multiple demes. Theoretical Population Biology, 79(3):97–101.

Priklopil, T. (2012). On invasion boundaries and the unprotected coexistence of two strategies. Journal of mathematical biology, 64:1137–1156.

Priklopil, T. and Lehmann, L. (2020). Invasion implies substitution in ecological communities with class-structured populations. Theoretical Population Biology, 134:36–52.

Priklopil, T. and Lehmann, L. (2021). Metacommunities, fitness and gradual evolution. Theoretical population biology, 142:12–35.

Priklopil, T. and Lehmann, L. (2024). On the interpretation of the operation of natural selection in class-structured populations. The American Naturalist, 203(2):292–304.

Rousset, F. (2004). Genetic Structure and Selection in Subdivided Populations. Princeton University Press, Princeton, NJ.

Slatkin, M. (1987). Gene flow and the geographic structure of natural populations. Science, 236(4803):787–792.

Smith, J. M. (1970). Genetic polymorphism in a varied environment. The American Naturalist, 104(939):487–490.

Stewart, G. W. (2001). Matrix Algorithms: Volume II: Eigensystems. SIAM.

Svardal, H., Rueffler, C., and Hermisson, J. (2015). A general condition for adaptive genetic polymorphism in temporally and spatially heterogeneous environments. Theoretical Population Biology, 99:76–97.

Tomasini, M. and Peischl, S. (2018). Establishment of locally adapted mutations under divergent selection. Genetics, 209(3):885–895.

Tuljapurkar, S. (1989). An uncertain life: demography in random environments. Theoretical Population Biology, 35:227–94.

Wilkinson, J. H. (1965). The algebraic eigenvalue problem, volume 662. Oxford Clarendon.

Wright, S. (1931). Evolution in mendelian populations. Genetics, 16:97–159.

Yeaman, S. (2015). Local adaptation by alleles of small effect. The American Naturalist, 186(S1):S74–S89.

Yeaman, S. and Otto, S. P. (2011). Establishment and maintenance of adaptive genetic divergence under migration, selection, and drift. Evolution, 65(7):2123–2129.

